# Neuronal feedback loop in the suprachiasmatic nucleus generates robust circadian rhythms

**DOI:** 10.1101/2025.05.21.655394

**Authors:** Mohan Wang, Yusuke Tsuno, Yubo Peng, Ayako Matsui, Takashi Maejima, Michihiro Mieda

## Abstract

The central circadian clock of the suprachiasmatic nucleus (SCN) is a network consisting of various types of neurons and glial cells, but its mechanism remains elusive. Here, by monitoring cellular calcium rhythms in vivo in vasoactive intestinal polypeptide (VIP)-deficient mice, we show that arginine vasopressin (AVP) neurons intrinsically oscillate with a short period and that VIP amplifies and delays the AVP neuronal rhythm to lengthen its period to ∼24 hours. Consistently, the circadian behavior period is shortened and lengthened by VIP receptor disruption and neurotransmission blockade of AVP neurons, respectively. VIP and other neurons occasionally exhibit weak, unstable, long-period calcium rhythms only when AVP neuronal oscillation is attenuated. Together with our previous finding that AVP neurons act as the primary pacesetter cells of the SCN ensemble rhythm, our results indicate that the feedback neuronal circuit of AVP cellular oscillator and VIP peptide signaling is crucial for the robust circadian rhythm.

## Introduction

The hypothalamic suprachiasmatic nucleus (SCN) acts as the central circadian pacemaker in mammals, orchestrating multiple circadian biological rhythms of behavior and physiological function according to the environmental light/dark (LD) cues conveyed from the eye.^1–4^ The SCN contains ∼20,000 cells, most of which can generate autonomous circadian oscillations. In individual cells, the circadian rhythm is driven by the autoregulatory transcription-translation feedback loop (TTFL) consisting of clock genes such as *Per1/2/3*, *Cry1/2*, *Bmal1*, and *Clock*.^1,2^ Notably, the TTFL is not unique to SCN cells and is common to almost all peripheral cells.^2,5^ Rather, the SCN constructs a network of many TTFL-driven cellular clocks to generate a highly robust and coherent circadian rhythm as the central clock, which entrains a large number of peripheral clocks throughout the body and is strikingly resistant to genetic and environmental perturbations.^1–4^ Thus, intercellular communication among SCN clock cells through the neuronal and diffusible network distinguishes the SCN from the peripheral clocks. However, the network mechanism underlying the SCN central clock remains unclear. The SCN is a heterogeneous structure composed of multiple types of neurons.^1–4^ Most SCN neurons are GABAergic, and several populations of these GABAergic neurons also co-express neuropeptides. These include arginine vasopressin (AVP)-producing neurons located predominantly in the dorsomedial part (the SCN shell) and vasoactive intestinal polypeptide (VIP)-producing neurons in the ventrolateral part (the SCN core). VIP is the most important contributor to the maintenance and synchrony of SCN cellular clocks with varying periods.^6,7^ Therefore, mice deficient in VIP (*Vip^−/−^*) or its receptor VPAC_2_ (*Vipr2^−/−^*) demonstrate a drastic attenuation of the free-running rhythm in constant darkness (DD) with a shortened period or unstable multiple periods. In the latter, the free-running period changes suddenly or two periods emerge simultaneously.^6,8–10^ VIP neurons receive direct projections from the retina, are involved in the photoentrainment, and send dense projections dorsally to the SCN shell.^1,9,11^ However, the precise mechanisms by which VIP regulates SCN circuit remain incompletely understood.

On the other hand, our studies and those of others have suggested that the TTFL-driven cellular clocks in AVP neurons may play a dominant role over those of other SCN neurons in the generation of circadian oscillation by the SCN network, regulating the circadian oscillation of other cells and setting the SCN ensemble period.^3^ Thus, AVP neuron-specific disruption of the TTFL by deleting *Bmal1* (*Avp-Bmal1^−/−^* mice) results in an unstable, attenuated free-running rhythm with a lengthened period and the activity time (the interval between the onset and offset of locomotor activity).^12–16^ Furthermore, the cellular period length of AVP neurons appears to be the primary determinant of the SCN ensemble period.^16,17^ In contrast, similar genetic manipulations of disrupting the TTFL or altering the cellular period in VIP neurons have little effect on the amplitude and period of the free-running behavior rhythm,^13–16,18^ suggesting that VIP peptide is critical but the TTFL in VIP neurons may be dispensable for the circadian timekeeping. Besides, our previous studies indicated the importance of in vivo analysis of SCN network dynamics.^16,17^

Thus, there is a functional differentiation between the shell and the core within the SCN network. However, how the two integrate to generate the circadian rhythm of the SCN ensemble is still unclear. In the current study, we investigated the regulation of AVP neuronal circadian oscillation by VIP in vivo. To this end, we applied dual-color fiber photometry recording, which allows us to monitor the cellular circadian rhythms from two types of SCN neurons simultaneously, to VIP-deficient mice. We then analyzed the circadian behavior rhythms of multiple mouse strains that had various impairments in the communication between VIP and AVP neurons. Our results reveal a feedback loop neuronal circuit of these two types of neurons that may be the essential network mechanism of the central circadian pacemaker of the SCN.

## Results

### SCN AVP neurons maintain a stable Ca^2+^ rhythm with a short period and reduced amplitude in VIP-deficient mice

To elucidate the mechanism of how VIP regulates the cellular clocks of AVP neurons, we first aimed to compare the circadian rhythm of intracellular Ca^2+^ in SCN AVP neurons between control and VIP-deficient mice. Intracellular Ca^2+^ rhythms are a nice measure of cellular circadian rhythms in vivo that can be recorded in a neuron type-specific manner using genetically encoded calcium sensors and fiber photometry. We used the *Vip-tTA* knock-in mice (*Vip^tTA/tTA^*), which are equivalent to *Vip* knock-out mice and exhibit a spectrum of phenotypes such as simple short-period or complicated multiple-period behavioral rhythms.^10^ In addition, their specific expression of tTA in VIP neurons allowed us to record Ca^2+^ rhythms in SCN AVP and VIP neurons (AVP- and VIP-Ca^2+^) separately and simultaneously when combined with hemizygous *Avp-Cre* BAC transgenic mice.^12^ Thus, we generated VIP-deficient *Avp-Cre; Vip^tTA/tTA^* mice and heterozygous control *Avp-Cre; Vip^wt/tTA^* mice and then targeted the expression of green (jGCaMP7s)^19^ and red (jRGECO1a)^20^ fluorescent Ca^2+^ sensor proteins in SCN AVP and VIP neurons, respectively, by simultaneous injection of Cre- and tTA-dependent AAV vectors (AAV-*CAG-FLEX-jGCaMP7s* and AAV-*TRE-jRGECO1a*) (Figures 1A and 1B). In vivo dual-color fiber photometry revealed robust, temporally coordinated AVP- and VIP-Ca^2+^ rhythms in control mice in both light/dark (LD) and dark/dark (DD), higher during the (subjective) day with a slightly earlier rise in AVP-Ca^2+^ (Figures 1C, 1D and S2A). In VIP-deficient mice, the VIP-Ca^2+^ rhythm was comparable to that of controls in LD but rapidly attenuated in DD, even when the locomotor activity still showed rhythmicity. In contrast, the amplitude of the AVP-Ca^2+^ rhythm was substantially reduced in both LD and DD, but it remained stable in DD (Figures 1C-1F and S1). Consistent with the changes in the behavior rhythm, the AVP-Ca^2+^ rhythm showed a phase advance in LD (midpoint phase: control, ZT 4.11 ± 0.42; VIP-deficient, ZT 21.60 ± 0.84, p < 0.001) and a shortened period in DD (control, ∼23.7h; VIP-deficient, ∼22.5 h, Figures 1C-1E, S2B and S2C, p < 0.05). Notably, the latter was almost identical to the short free-running period of the behavior rhythm, even when mouse behavior changed to a long-period rhythm later in DD.

**Figure 1.**
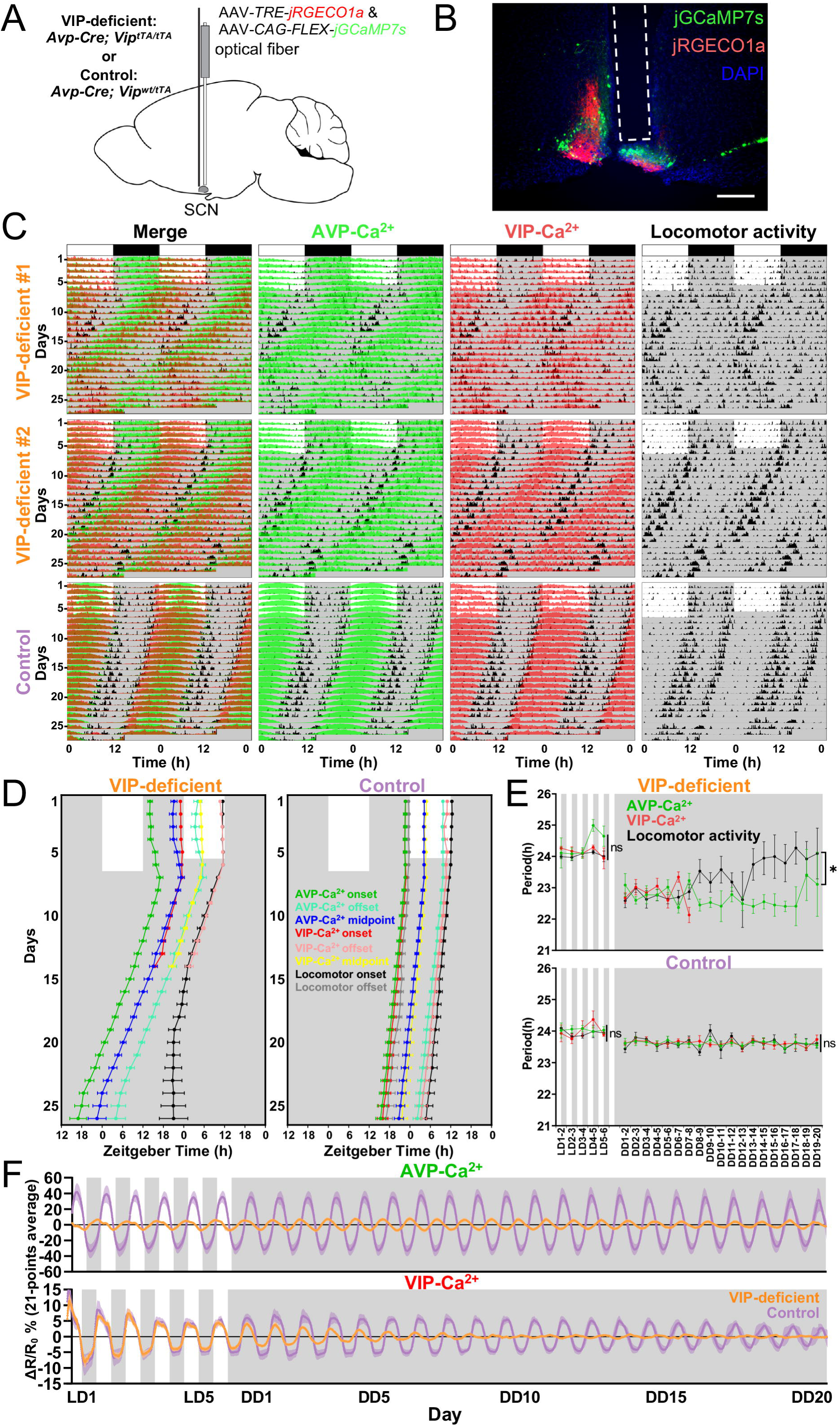
SCN AVP neurons maintain a stable Ca^2+^ rhythm with a short period and reduced amplitude in VIP-deficient mice. **(A)** Schematic diagram of viral vector (AAV-*CAG-FLEX-jGCaMP7s* and AAV-*TRE-jRGECO1a*) injection and optical fiber implantation at the SCN in control (*Avp-Cre; Vip^wt/tTA^*) or VIP-deficient (*Avp-Cre; Vip^tTA/tTA^*) mice for fiber photometry recording. **(B)** A representative coronal SCN section of mice with jGCaMP7s expression in AVP neurons and jRGECO1a in VIP neurons. A white dotted square shows the estimated position of the implanted optical fiber. Green, jGCaMP7s; red, jRGECO1a; blue, DAPI. Scale bar, 200 μm. **(C)** Representative plots of the in vivo jGCaMP7s signal of AVP neurons (AVP-Ca^2+^, green) and jRGECO1a signal of VIP neurons (VIP-Ca^2+^, red) overlaid with the locomotor activity (home-cage activity) (black) in actograms. Ca^2+^ signals are shown with normalization in each row. VIP-deficient and control mice were initially housed in 12:12-h LD and then in DD. Gray shading indicates the time when the lights were off. **(D)** Plots of locomotor activity onset (black), activity offset (gray), AVP-Ca^2+^ onset (green), AVP-Ca^2+^ offset (cyan), AVP-Ca^2+^ midpoint (blue), VIP-Ca^2+^ onset (red), VIP-Ca^2+^ offset (light red), and VIP-Ca^2+^ midpoint (yellow) in VIP-deficient and control mice. The activity offset and some parts of VIP-Ca^2+^ values are unmeasurable in VIP-deficient mice due to their attenuated rhythmicity. **(E)** Day-by-day changes in the periods of AVP-Ca^2+^ (green), VIP-Ca^2+^ (red), and the locomotor activity (black) rhythms in LD and DD. *p < 0.05 by two-way repeated measures ANOVA. ns, not significant. **(F)** Time courses of AVP-Ca^2+^ and VIP-Ca^2+^ rhythms for 26 days (6 days in LD, 20 days in DD). Detrended and smoothened data from control (lavender) and VIP-deficient (orange) mice are shown. Values are mean ± SEM. n=5 for control; n=8 for VIP-deficient mice **(C-F)**.

These results suggest that AVP neurons intrinsically oscillate with a short period and that VIP enhances and delays the oscillation of AVP neurons to make their period closer to 24 h. In turn, the timing of VIP release may be regulated by the cellular rhythm of AVP neurons.

### VIP-Ca^2+^ changes to a weak, long-period rhythm at the late stage in DD

In VIP-deficient mice, the AVP-Ca^2+^ rhythm was almost correlated with the short-period behavior rhythm in DD, suggesting that AVP neurons were responsible for it. As a result, in more than half of VIP-deficient mice, the AVP-Ca^2+^ rhythm dissociated from the behavior rhythm in the later stage of DD, which became a long-period rhythm. The VIP-Ca^2+^ rhythm in actograms initially followed the short-period AVP-Ca^2+^ and behavior rhythms but gradually disappeared and became very weak and noisy in DD (Figures 1C-1F). Considering the multiple periodicities of the behavior rhythm in VIP-deficient mice,^6,8–10^ we performed periodogram analyses of Ca^2+^ rhythms throughout three consecutive weeks in DD to detect weak rhythms with different periods. The VIP-Ca^2+^ rhythm demonstrated two clear period peaks in most VIP-defficient mice, namely a primary short period (22.67 h) and a secondary long period (24.33 h), whereas AVP-Ca^2+^ rhythm showed only one short period (22.67 h) (Figure 2A, left and Figure S1). In contrast, both AVP- and VIP-Ca^2+^ rhythms of control mice presented a robust single period within the normal range (23.67 h), which matched that of the behavior rhythm (Figure 2A, right and Figure S2A).

**Figure 2.**
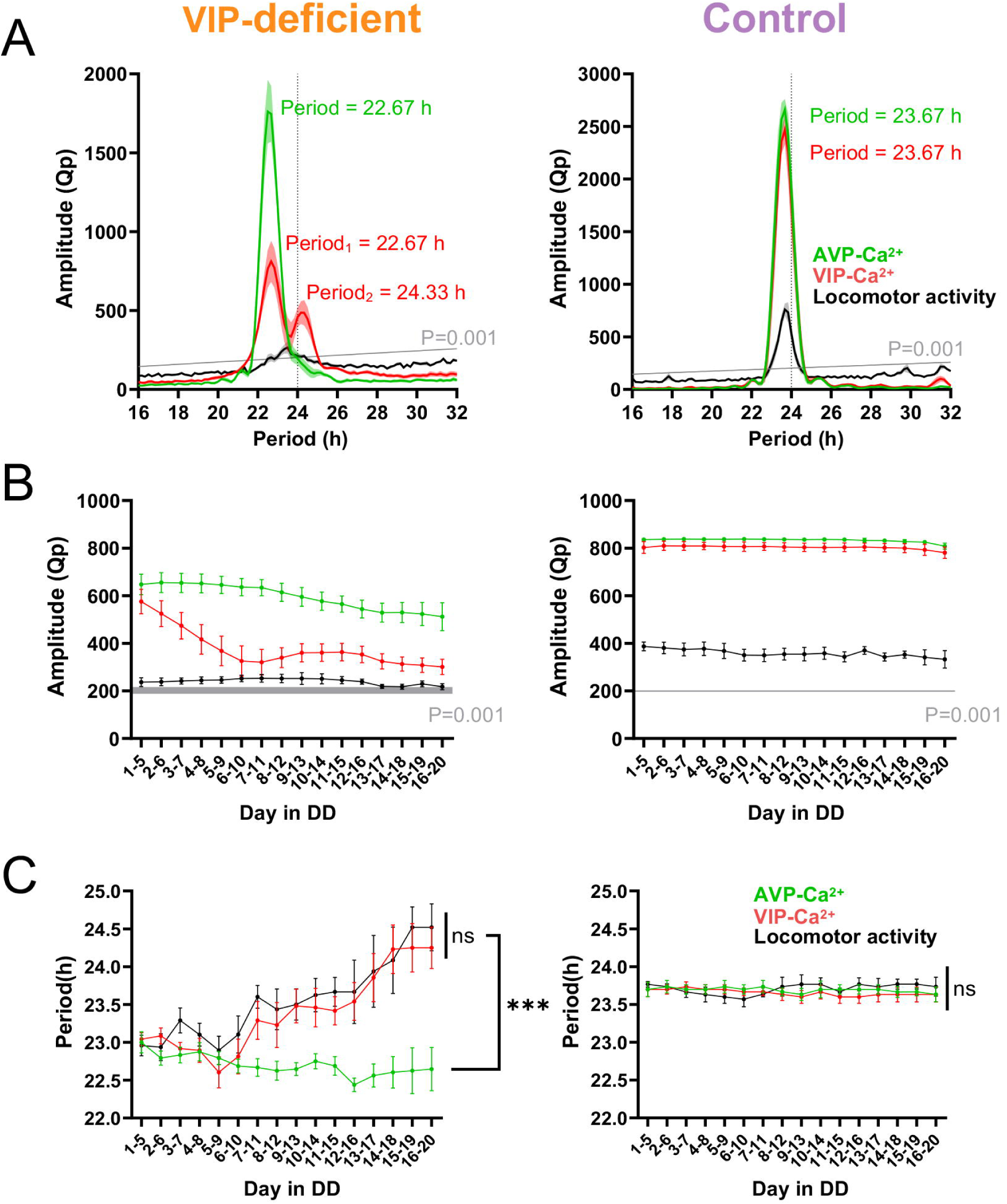
VIP-Ca^2+^ changes to a weak, long-period rhythm at the late stage in DD in VIP-deficient mice. **(A)** Averaged periodograms of detrended AVP-Ca^2+^ (green), VIP-Ca^2+^ (red) signals, and the locomotor activity (black) of VIP-deficient and control mice for days 1–20 in DD. Gray line, significance level; black dotted line, 24 h. **(B and C)** Time courses of amplitudes (Qp) **(B)** and periods **(C)** of AVP-Ca^2+^ (green), VIP-Ca^2+^ (red), and the locomotor activity (black) rhythms in DD by sequential periodogram analyses with shifting 5-day windows. Values are mean ± SEM. n = 5 for control; n = 8 for VIP-deficient mice. ***p < 0.001 by two-way repeated measures ANOVA; ns, not significant.

To find out at which stage of DD the secondary long period occurred, we next calculated the time courses of amplitudes (Qp) and periods of AVP-Ca^2+^, VIP-Ca^2+^, and behavior rhythms by sequential periodogram analyses with shifting 5-day windows. The peak Qp values of the VIP-Ca^2+^ rhythm decreased continuously during the first 7-8 days (early stage) and then sustained at a relatively low value until the end of the recording (late stage) (Figure 2B, left). At the early stage, there was no significant difference in the period length between AVP-Ca^2+^, VIP-Ca^2+^, and behavior rhythms (Figure 2C, left). At the late stage, however, the period of the VIP-Ca^2+^ rhythm was lengthened close to the long period of the behavior rhythm (VIP-Ca^2+^ vs. Locomotor activity, p = 0.46), whereas the AVP-Ca^2+^ period remained short (AVP-Ca^2+^ vs. VIP-Ca^2+^, p < 0.001; AVP-Ca^2+^ vs. Locomotor activity, p < 0.001, Figure 2C, left and Figure S1). In control mice, all periods of Ca^2+^ and behavior rhythms were stable and consistent (Figure 2B, right; Figure 2C, right and Figure S2A). These results suggest that VIP neurons may be a subordinate slow oscillator and exhibit a weak, long-period rhythm that correlates with the behavior rhythm in vivo when the AVP cellular rhythm is attenuated by VIP deficiency.

### Ca^2+^ rhythm of CCK neurons in VIP-deficient mice rapidly attenuates in DD

Next, we aimed to know how unique the consistently short-period AVP-Ca^2+^ and the late-onset, weak, long-period VIP-Ca^2+^ rhythms are. In particular, CCK neurons enriched in the anterior shell region of the SCN were of interest as another candidate for a long-period oscillator, because they reportedly play an important role in the behavior under a long-day photoperiod and their ablation slightly shortened the behavioral free-running period in DD.^21^ To test this possibility, we introduced a *Cck-ires-Cre* allele^22^ to VIP-deficient (*Cck^wt/ires-Cre^; Vip^tTA/tTA^*) and control (*Cck^wt/ires-Cre^; Vip^wt/tTA^*) mice and labeled CCK and VIP neurons with jGCaMP7s and jRGECO1a, respectively, by injecting AAV vectors into the SCN (Figures 3A and 3B). Our fiber photometry recording successfully recapitulated the reported CCK-Ca^2+^ rhythm in control mice, with its peak in the second half of the (subjective) day (Figures 3C and 3D).^21^ In VIP-deficient mice, the CCK-Ca^2+^ rhythm was robust in LD but rapidly attenuated in DD, as did VIP-Ca^2+^ (Figures 3C, 3D, and S3). However, in contrast to the VIP-Ca^2+^ rhythm, the periodogram analysis of the CCK-Ca^2+^ rhythm for 3-week recordings in DD showed only one period peak (22.00 h), which was even slightly shorter than that of AVP-Ca^2+^ (22.67 h), but with a broad distribution of significant period lengths on the long-period side up to ∼24.3 h, i.e., the secondary peak of the VIP-Ca^2+^ period (Figure 3E). Sequential periodogram analysis revealed that the CCK-Ca^2+^ rhythm gradually reduced its amplitude (Qp) and maintained a short period on average, but with a large variability. In contrast, in control mice, CCK-Ca^2+^, VIP-Ca^2+^, and behavioral periods were stable and consistent (23.50 h) (Figures 3E-3G). These results suggest that the CCK-Ca^2+^ rhythm is weak and unstable and has more heterogeneous periods in VIP deficiency. Such heterogeneity may partly reflect that CCK neurons contain at least two subpopulations, one overlapping with a small subpopulation of AVP neurons.^23,24^

**Figure 3.**
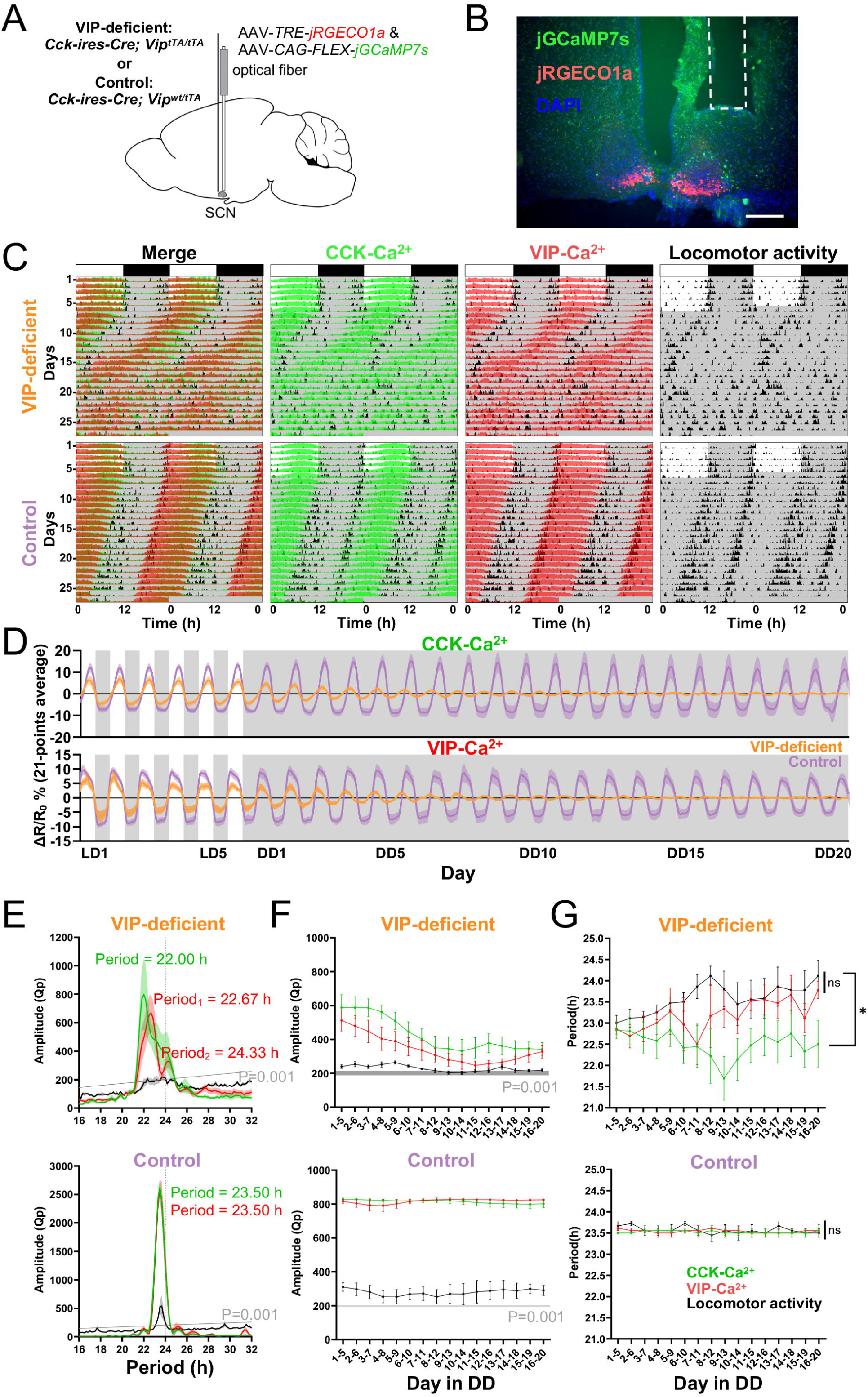
CCK-Ca^2+^ rhythm in VIP-deficient mice rapidly attenuates in DD. **(A)** Schematic diagram of viral vector (AAV-*CAG-FLEX-jGCaMP7s* and AAV-TRE-jRGECO1a) injection and optical fiber implantation at the SCN in control (*Cck^wt/ires-Cre^; Vip^wt/tTA^*) or VIP-deficient mice (*Cck^wt/ires-Cre^; Vip^tTA/tTA^*) for fiber photometry recording. **(B)** A representative coronal SCN section of mice with jGCaMP7s expression in CCK neurons and jRGECO1a in VIP neurons. A white dotted square shows the estimated position of the implanted optical fiber. Green, jGCaMP7s; red, jRGECO1a; blue, DAPI. Scale bar, 200 μm. **(C)** Representative plots of the in vivo jGCaMP7s signal of CCK neurons (CCK-Ca^2+^, green) and jRGECO1a signal of VIP neurons (VIP-Ca^2+^, red) overlaid with the locomotor activity (black) in actograms. Ca^2+^ signals are shown with normalization in each row. VIP-deficient and control mice were initially housed in 12:12-h LD and then in DD. Gray shading indicates the time when the light was off. **(D)** Time courses of CCK-Ca^2+^ and VIP-Ca^2+^ rhythms for 26 days (6 days in LD, 20 days in DD). Detrended and smoothened data from control (lavender) or VIP-deficient (orange) mice are shown. **(E)** Averaged periodogram of detrended CCK-Ca^2+^ (green), VIP-Ca^2+^ (red) signals and the locomotor activity (black) of VIP-deficient and control mice for days 1–20 in DD. Gray line, significance level; black dotted line, 24 h. **(F and G)** Time courses of amplitudes (Qp) **(F)** and periods **(G)** of CCK-Ca^2+^ (green), VIP-Ca^2+^ (red), and the locomotor activity (black) rhythms in DD by sequential periodogram analyses with shifting 5-day windows. *p < 0.05 by two-way repeated measures ANOVA; ns, not significant. Values are mean ± SEM. n = 3 for control; n = 6 for VIP-deficient mice **(D-F)**.

### Ablation of VIP neurons in adulthood causes a simple short-period behavior rhythm

VIP-deficient *Vip^tTA/tTA^* mice free-ran initially with a short period in DD, but about 60 % of the mice changed the free-running rhythm to a long period after 1∼2 weeks (Figures S1 and S3). This observation was consistent with the previous reports of *Vip* and *Vipr2* knockout mice.^6,8,9^ On the other hand, Mazuski et al. reported that mice in which VIP neurons were ablated in adulthood by Caspase 3-mediated apoptosis retained the wheel running rhythm but with a shortened period (∼22.7 h).^25^ We also performed a similar experiment to measure the locomotor activity (home cage activity) rhythm by injecting an AAV vector expressing activated Caspase 3 in a Cre-dependent manner (AAV-*CAG-FLEX-taCasp3-TEVp*) into the SCN of *Vip^wt/ires-Cre^* mice. We obtained a similar result, i.e., a shortened free-running period and reduced amplitude of the rhythm (Figure S4). These results suggest that VIP neurons contribute to circadian timekeeping by lengthening the circadian period and promoting the rhythmicity in adults. Some compensatory mechanisms may cause multiple periods in VIP-deficient mice during development.

### Dysfunction of VPAC_2_ in AVP neurons acutely shortens the free-running period

We next investigated the importance of VIP/VPAC_2_ signaling in AVP neurons for the pacesetting of the SCN network. We used in vivo genome editing to selectively disrupt the *Vipr2* gene in AVP neurons. First, we crossed *Avp-Cre* mice with *Rosa26-LSL-SpCas9* mice, which express SpCas9 Cre-dependently. We then injected AAV vectors expressing gRNAs targeting the *Vipr2* (AAV-*U6-gVipr2-EF1α-DIO-mCherry*, *Avp-Vipr2^−/−^* mice) or control gRNAs (AAV-*U6-gControl-EF1α-DIO-mCherry*) into the SCN of *Avp-Cre; Rosa26-LSL-SpCas9* mice (Figures 4A and 4B).

**Figure 4.**
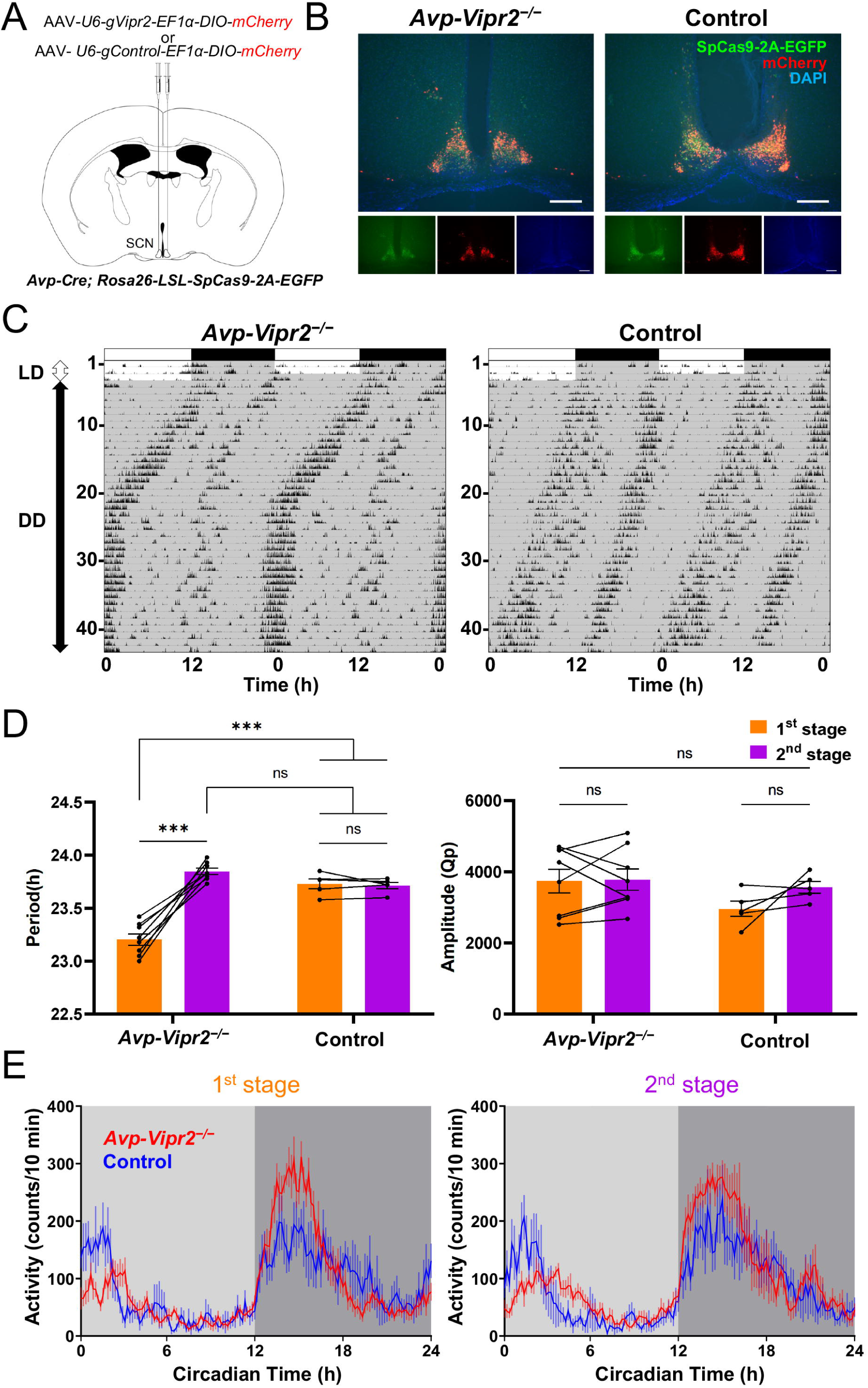
VPAC_2_ dysfunction in AVP neurons acutely shortens the free-running period. **(A)** Schematic diagram showing the injection of AAV-*U6-gVipr2-EF1α-DIO-mCherry* or AAV-*U6-gControl-EF1α-DIO-mCherry* into the SCN of *Avp-Cre; Rosa26-LSL-SpCas9-2A-EGFP* mice. **(B)** Representative coronal SCN sections prepared from *Avp-Cre; Rosa26-LSL-SpCas9* mice with a targeted AAV injection. Scale bars, 200 mm; green, SpCas9-2A-EGFP; red, mCherry; blue, DAPI). **(C)** Representative locomotor activity actograms of *Avp-Vipr2^−/−^* and control mice. Recordings were started immediately after the AAV injection to track the course of behavior change. Mice were initially housed in LD for 3 days and then in DD. Gray shading indicates the time when the light was off. **(D)** The free-running period and amplitude (Qp values) of the locomotor activity rhythm in DD. Because *Avp-Vipr2^−/−^* obviously altered the free-running period in the middle of DD, periodogram analysis was performed separately before (1st stage) and after (2nd stage) the transition point for 10 days when the periods were stable. Values are mean ± SEM; n = 8 for *Avp-Vipr2^−/−^*, n = 6 for control mice. ***p < 0.001 by two-way repeated measures ANOVA; ns, not significant. **(E)** Averaged daily profiles of the locomotor activity of 1st and 2nd stages in DD.

*Avp-Vipr2^−/−^* mice free-ran with a period significantly shorter than controls when released into DD (23.21 ± 0.05 h vs. 23.64 ± 0.10 h, p < 0.001) (Figures 4C, 4D, and S5). Surprisingly, however, after a while, *Avp-Vipr2^−/−^* mice suddenly changed their free-running period to the one comparable to controls (23.85 ± 0.03 h vs. 23.60 ± 0.11 h, p = 0.18). This observation led us to start recording locomotor activity immediately after the AAV injection in most of the mice examined, with recordings in LD only for 3 days before DD (Figure S5). Strikingly, all *Avp-Vipr2^−/−^* mice exhibited changes in the free-running period around 21 days after AAV injection. Control mice never showed such a sudden change in the free-running period (Figure 4C, 4D, and S5). These results indicate that disruption of VIP/VPAC_2_ signaling in AVP neurons in adulthood acutely shortens the free-running period due to the intrinsic short period of AVP neuronal clocks in the absence of phase-delaying regulation by VIP. On the other hand, the period shortening in *Avp-Vipr2^−/−^* mice was less than in VIP-deficient mice. Moreover, the shortening was abolished ∼3 weeks after the blockade of the VIP → AVP neuron axis. These observations suggest indirect pathways in the regulation of AVP neurons by VIP neurons that may be involved in the compensatory recovery of the standard period in *Avp-Vipr2^−/−^* mice. Such tight regulation of short-period AVP neuronal oscillation may imply its fundamental importance in the circadian pacemaking of the SCN network.

### Disrupting TTFLs in AVP neurons leaves an attenuated VIP-Ca^2+^ rhythm with a long period

We previously reported that *Bmal1* deletion specific to AVP neurons (*Avp-Cre; Bmal1^flox/flox^* = *Avp-Bmal1^−/−^* mice) leaves an attenuated circadian behavior rhythm with a lengthening of the free-running period and the activity time.^12^ In contrast, mice become completely arrhythmic when *Bmal1* is deleted in all SCN neurons using forebrain-specific *CamKII-Cre* driver mice.^16,26^ These results may be explained by the weak circadian oscillators of SCN non-AVP neurons with a long period as a population in vivo, including VIP and CCK neurons as described above. To test this possibility, we recorded AVP-Ca^2+^ and VIP-Ca^2+^ rhythms in *Avp-Bmal1^−/−^* mice. To do so, we generated *Avp-Bmal1^−/−^; Vip^wt/tTA^* mice and injected AAV vectors (AAV-*CAG-FLEX-jGCaMP7s* and AAV-*TRE-jRGECO1a*) into the SCN (Figures 5A and 5B). As expected, the AVP-Ca^2+^ exhibited little circadian rhythmicity, although this might be due to low jGCaMP7s expression in TTFL-deficient AVP neurons (Figures 5C, 5E, and S6). Notably, the VIP-Ca^2+^ retained a rhythm with a long period (24.17 h) that is consistent with the period of the behavior rhythm (24.17 h) in DD (Figures 5E and S6B). The duration of high VIP-Ca^2+^ in the subjective day was compressed according to the prolongation of the activity time of the behavior rhythm (Figures 5C, 5D, and S6A). Consistent with the previous report that a small number of *Avp-Bmal1^−/−^*mice show multiple periods,^12^ one out of four *Avp-Bmal1^−/−^; Vip^wt/tTA^* mice demonstrated a similar multiple period phenotype. In this particular mouse, the VIP-Ca^2+^ rhythm was significant but largely attenuated (Figure S6A, mouse #4). These results support the idea that an attenuated oscillation of non-AVP neurons with a long period on average manifests in the circadian behavior rhythm.

**Figure 5.**
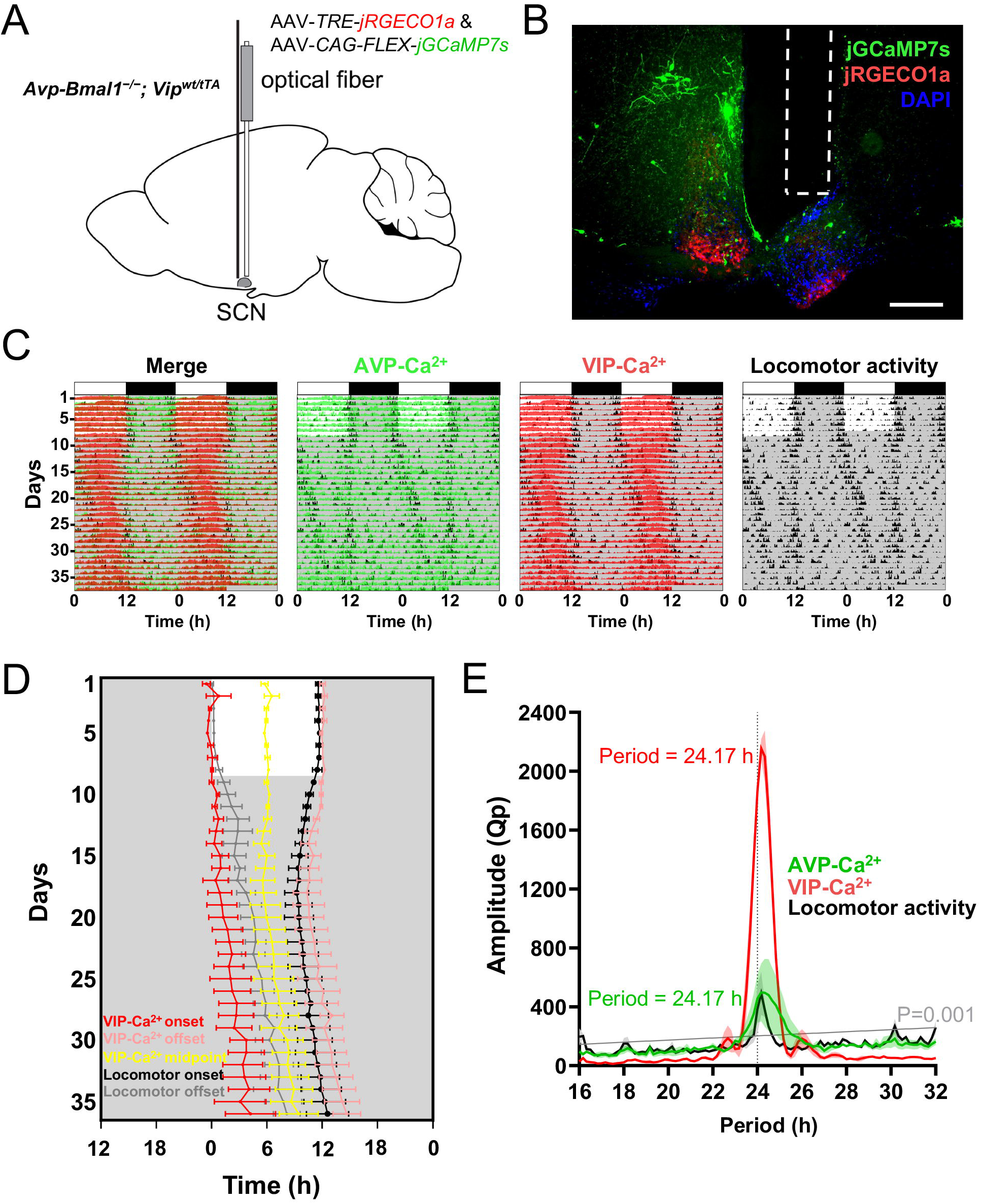
*Avp-Bmal1^−/−^* mice show the VIP-Ca^2+^ rhythm with a long period. **(A)** Schematic diagram of viral vector (AAV-*CAG-FLEX-jGCaMP7s* and AAV-*TRE-jRGECO1a*) injection and optical fiber implantation at the SCN in *Avp-Bmal1^−/−^; Vip^wt/tTA^*mice for fiber photometry recording. **(B)** A representative coronal SCN section of mice with jGCaMP7s expression in AVP neurons and jRGECO1a in VIP neurons. A white dotted square shows the estimated position of the implanted optical fiber. Green, jGCaMP7s; red, jRGECO1a; blue, DAPI. Scale bar, 200 μm. **(C)** Representative plots of the in vivo jGCaMP7s signal of AVP neurons (AVP-Ca^2+^, green) and jRGECO1a signal of VIP neurons (VIP-Ca^2+^, red) overlaid with the locomotor activity (black) in actograms. Ca^2+^ signals are shown with normalization in each row. Mice were initially housed in LD and then in DD. Gray shading indicates the time when the lights were off. **(D)** Plots of locomotor activity onset (black), activity offset (gray), VIP-Ca^2+^ onset (red), VIP-Ca^2+^ offset (light red), and VIP-Ca^2+^ midpoint (yellow). **(E)** Averaged periodogram of detrended AVP-Ca^2+^ (green), VIP-Ca^2+^ (red) signals, and the locomotor activity (black) of *Avp-Bmal1^−/−^; Vip^wt/tTA^* mice for the last 14 days in DD. Values are mean ± SEM. n = 3. Gray line, significance level; black dotted line, 24 h.

### Blocking neurotransmitter release from AVP neurons lengthens the free-running period

To further explore the evidence that AVP neurons act as the principal oscillator with an intrinsically short period in the SCN, we next examined the behavioral changes in mice in which the neurotransmitter release from SCN AVP neurons was blocked by AAV-mediated expression of tetanus toxin light chain (*Avp-TeNT* mice, AAV-*EF1α-DIO-GFP::TeNT* injected into the SCN of *Avp-Cre* mice) (Figures 6A and 6B). As expected, *Avp-TeNT* mice showed a longer free-running period (24.48 ± 0.06 h) in DD than control mice (23.29 ± 0.06 h, p < 0.001, Figures 6C and 6D). The amplitude of the locomotor activity rhythm, as measured by the Qp values of the periodogram, was not significantly different between groups (p = 0.64, Figure 6D). However, individual *Avp-TeNT* mice exhibited diverse daily patterns of locomotor activity that deviated significantly from the normal nocturnal and bimodal pattern with clear evening (E) and morning (M) locomotor activities (Figures 6C and S7A). These data further confirmed that AVP neurons intrinsically encode a short-period circadian rhythm.

**Figure 6.**
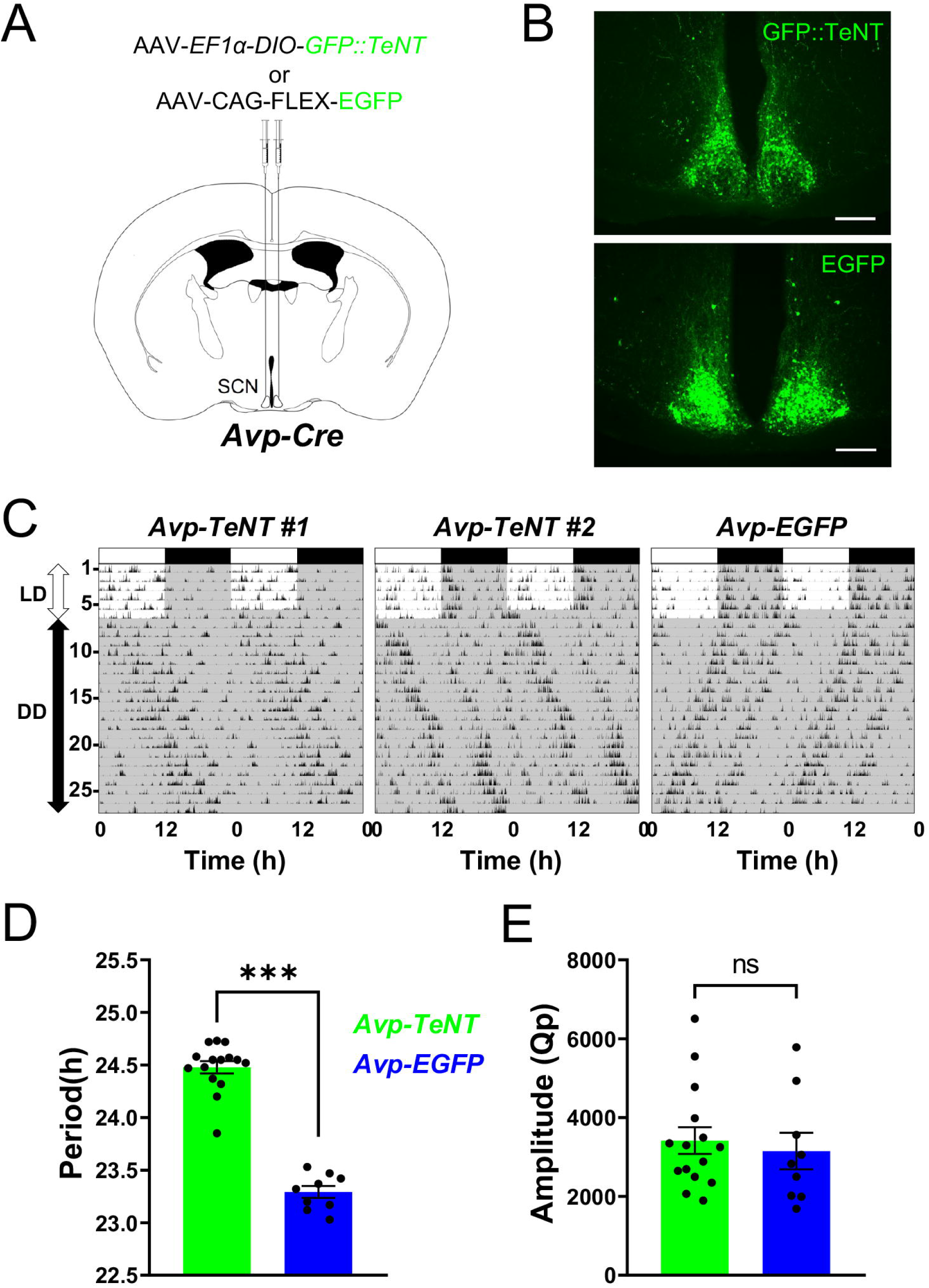
Blocking neurotransmitter release from AVP neurons lengthens the free-running period. **(A)** Schematic diagram showing the injection of AAV-*EF1α-DIO-GFP::TeNT* or AAV-*CAG-FLEX-EGFP* into the SCN of *Avp-Cre* mice. **(B)** Representative coronal SCN sections of mice with GFP::TeNT (immunostained with anti-GFP antibody) or EGFP expression. Scale bar, 200 μm. **(C)** Representative locomotor activity actograms of *Avp-TeNT* or *Avp-EGFP* (control) mice. Animals were initially housed in LD and then in DD. Gray shading indicates the time when the light was off. **(D and E)** The free-running period and amplitude (Qp values) of the locomotor activity rhythm in DD. Values are mean ± SEM; n = 15 for *Avp-TeNT*, n = 9 for *Avp-EGFP* mice. ***p < 0.001 by two-tailed Welch t test; ns, not significant.

To our surprise, many *Avp-TeNT* mice showed varying degrees of diurnal locomotor activity rhythms in both LD and DD (Figure 6C). To understand the underlying cause, we followed the time course of the emergence of apparent diurnal rhythms in LD after the onset of TeNT expression. Interestingly, TeNT expression in AVP neurons rapidly altered the coupling of E and M locomotor activities (Figure S7A). Namely, *Avp-TeNT* mice quickly exhibited a prolonged activity time, specifically manifested as a slight advancement in locomotor activity onset (i.e., E activity) and a significant delay in locomotor activity offset (i.e., M activity). A new steady state of the M-E (M is closer to the following E than to the preceding E) phase relation, rather than the normal E-M phase relation, generated an apparent diurnal behavior rhythm in some mice that persisted in DD. Such features can be regarded as a more severe phenotype of the lengthened activity time observed in *Avp-Bmal1^−/−^* and *Avp-Vgat^−/−^*mice.^12,27^

## Discussion

### In vivo fiber photometry recording is a powerful method to investigate the SCN network dynamics

First, the fiber photometry recordings in this study measured the Ca^2+^ rhythms of neuronal populations defined by the expression of a particular neuropeptide (i.e., AVP, VIP, or CCK).^11,16,21,27,28^ Therefore, the estimated period, phase, and amplitude values may reflect the averages of many neurons with various circadian oscillations within each population. In addition, each of these neuronal populations still contains molecularly heterogeneous, multiple subpopulations that may have different rhythms and functions.^23,24^ Nevertheless, this method is so far the only one that allows long-term recordings of SCN dynamics in genetically defined neuronal populations from freely behaving mice. Furthermore, the dual-color fiber photometry recording in this study is the first report of monitoring Ca^2+^ rhythms from two types of neurons simultaneously in vivo. As discussed below, the results obtained here provided sufficient information to gain insight into the network mechanism underlying the central clock of the SCN.

### The SCN neuronal feedback loop (SNFL)

Several studies have indicated that multiple circadian clocks with different period lengths in the SCN lose coupling in the absence of VIP signaling, resulting in the multiple-period behavior rhythm.^6–10^ In this study, we showed that, in such a condition, the AVP-Ca^2+^ rhythm was substantially phase-advanced in LD and continued to oscillate stably in DD with a short period following the dominant short-period behavior rhythm, although its amplitude was significantly reduced. In contrast, the VIP-Ca^2+^ rhythm, as well as the CCK-Ca^2+^ rhythm, was normal in LD but rapidly disappeared in DD. Taken together with our previous findings that AVP neurons act as the primary circadian pacesetter cells in vivo,^3,12,16,17^ we propose a feedback loop neural circuit of AVP and VIP neurons within the SCN as a critical network machinery of the mammalian central clock. Here, AVP neurons are the principal oscillatory part of the SCN and regulate the circadian rhythms of VIP and other neurons. Crucially, they are a fast oscillator with an intrinsically short period. VIP peptide, released rhythmically, acts on AVP neurons, delaying the AVP neuronal rhythm to lengthen its period closer to 24 h and increase its amplitude. Therefore, in the absence of VIP, the AVP neuronal rhythm expresses an intrinsic short period, but reduces its amplitude and weakens its control over other multiple clocks in the SCN with various period lengths. Such impairments manifest as the short-period or the multiple-period behavior rhythms. This scheme fits well with the reported significant phase-delaying effect of VIP and VIP neuronal activation on the SCN and behavior rhythms.^29–32^

Approximately 40 % of *Vip^−/−^* or *Vipr2^−/−^* mice demonstrate an attenuated circadian behavior rhythm with a single short period.^6,8–10^ Even in *Vip^−/−^* or *Vipr2^−/−^* mice showing multiple periodicity, the dominant periods are often short.^6^ Indeed, ablation of VIP neurons in adult mice results in a single short period (Figure S4),^25^ suggesting that the multiple periodicity may result from a chronic lack of VIP signaling throughout the development. Furthermore, the absence of VPAC_2_ in AVP neurons significantly shortened the free-running period, underscoring the importance of the VIP neuron → AVP neuron peptidergic axis.^32^ Intriguingly, however, the period shortening caused by *Vipr2* mutation in AVP neurons was less than that caused by VIP deficiency and disappeared after ∼3 weeks. These results implicate indirect, redundant, and compensatory pathways in the VIP regulation of the AVP neuronal period to tightly control the SCN ensemble period. Thus, the SNFL, which modifies the intrinsically short-period oscillation of AVP neurons to set the SCN ensemble period close to 24 h, may be the most critical network mechanism of the mammalian central clock.

It remains unknown whether such a short-period rhythm of AVP neurons is inherent in individual neurons or arises from the in vivo condition. Several studies have reported that AVP neurons or AVP-rich regions of the SCN show shorter periods in organotypic slices.^13,33–37^ Alternatively, in vivo conditions of the SCN, such as the network among AVP neurons, the interaction between AVP and other neurons, and the ambient humoral conditions, may set the short period of the AVP neuronal population. Indeed, we previously reported that the ability of AVP neurons to set the SCN ensemble period in vivo is lost in slices.^16^

### Cellular substrates of the weak long-period behavior rhythm

Because AVP neurons appear responsible for the short-period behavior rhythm, we next questioned the identity of the slow oscillators responsible for the long-period behavior rhythm in VIP-deficient mice. CCK neurons were a good candidate because they have been reported to track the onset of circadian behavioral activities and to coordinate the circadian activities under long photoperiods. Moreover, ablation of CCK neurons shortened the free-running period in DD by 10∼20 minutes, although TeNT expression in these cells did not alter the period length.^21^ We initially expected the presumable slow oscillator to continue consistently and independently of the fast oscillator in DD. However, the CCK-Ca^2+^ rhythm behaved similarly to VIP neurons in VIP-deficient mice, initially following the fast oscillator and rapidly reducing its amplitude. These observations may rule out the possibility that CCK neurons act as the discrete slow circadian oscillator in DD. Instead, VIP neurons often recovered weak Ca^2+^ rhythms with a lengthened period comparable to the behavioral period at the late stage after the initial attenuation. Because these VIP neurons lack VIP peptide and thus have substantially attenuated effects on the other cells, their weak, slow Ca^2+^ rhythm may be a passive oscillation. The late-stage periods of the CCK-Ca^2+^ rhythm were more variable, ranging from a shorter period than AVP-Ca^2+^ to a longer period similar to VIP-Ca^2+^. Therefore, the late-onset, slow, weak Ca^2+^ rhythms may be a general feature of multiple types of SCN neurons, potentially reflecting something like a background oscillation or a default state of the SCN that may play a marginal role in the pacesetting of the intact SCN network. Indeed, these long-period Ca^2+^ rhythms appear only when the functions of AVP neurons are attenuated (e.g., VIP-deficient mice, *Avp-Bmal1^−/−^* mice, and *Avp-TeNT* mice). The slight shortening of the free-running period by ablating CCK neurons may reflect their contribution to the indirect pathway controlling the AVP neurons’ short-period oscillation by VIP, which was discussed above. This idea is supported by the fact that CCK neurons reportedly respond to the activation of VIP neurons and activate AVP neurons.^21^

What makes such a presumable background oscillation? In mammals, almost every cell throughout the body has a TTFL, as do all SCN cells, with widely varying period lengths.^1,2^ Therefore, it is not surprising that weak circadian rhythms with various periods emerge when the control of primary oscillators (AVP neurons) over the rest of the SCN cellular oscillators is attenuated. Indeed, TTFL disruption in all SCN neurons resulted in a complete arrhythmicity (*CaMKII2α-Bmal1^−/−^* mice).^16,26^ Why is the background oscillation period longer than 24 h? It is highly challenging to know the accurate intrinsic cellular periods of individual SCN neurons in vitro, as they seem to depend largely on culture conditions (e.g., slice or dispersal, density of dispersal) and measurement methods (e.g., clock gene reporters, Ca^2+^, firing rate). Nevertheless, not all, but many mouse studies using *Per1-Luc*, *Per2::Luc*, or GCaMP reporters suggested that the average of highly heterogeneous cellular periods is slightly longer than 24 h.^38–42^ Such a feature is common to peripheral tissues and even to fibroblast cultures.^40,43–45^ Although it is unclear whether the cellular periods differ between SCN neuron types in vitro, it is tempting to assume that AVP neurons have relatively shorter intrinsic periods while the rest of the cells have relatively longer periods on average. Because the regulation of the SCN network by AVP neurons attenuates ex vivo,^16^ the SCN slices may tend to show circadian periods longer than 24 h.

The firing rate rhythms of individual SCN neurons also demonstrate diverse period lengths with an average of around 23.5 h in neonatal dispersals or slices of normal mice.^6,46–50^ Notably, VIP- or VPAC_2_-deficiency further broadens their distribution, which is consistent with the emergence of multiple periodicities in the behavior rhythms.^6,50^

### Dual oscillator model: comparison with the fly’s central clock network

The dual oscillator model may provide an alternative explanation for our experimental results. Namely, the coupling of a fast oscillator composed of AVP neurons and a slow oscillator of another unidentified cell population determines the integrated ensemble period of the SCN network. However, we prefer the above-mentioned phase-delaying SNFL model for the following reasons. Our previous studies indicated that AVP neurons are the dominant pacesetter cells in the SCN network in vivo.^16^ This view is consistent with the current finding that AVP neurons are the most stable oscillators in VIP deficiency, although attenuated compared to the intact condition with VIP signaling. Based on the Ca^2+^ rhythms in VIP-deficient mice in DD, assume that the in vivo intrinsic period length of the dominant fast oscillator (i.e., AVP neurons) is ∼22.7 h and that of the presumable weak slow oscillator is ∼24.3 h. The free-running period of the locomotor activity rhythm of control mice is approximately ∼23.7 h. If the ensemble period is set according to a simple coupling of stronger fast and weaker slow oscillators, the ensemble period length should be much closer to that of the fast oscillator, which is not the case. Moreover, VIP-mediated coupling of these two oscillators should synergistically increase the amplitude of the ensemble oscillation in the intact mice. However, it would be unlikely that coupling two oscillators with different frequencies would greatly increase the amplitude of the integrated oscillation. Therefore, a simple coupling of two oscillators with different period lengths seems insufficient to explain our results.

The dual oscillator model was originally developed to account for several circadian properties of nocturnal rodents, including bimodal activity patterns and the ability to measure seasonal changes of day length.^51^ This model assumes two separate circadian oscillators, the morning (M) and evening (E) oscillators, which drive morning and evening locomotor activities, respectively, and are decelerated and accelerated by the external light. M and E oscillators are normally coupled to maintain a constant length of the activity time. In Drosophila, the M and E peaks of locomotor activity are differentially controlled by the pigment-dispersing factor (PDF)-positive small ventral lateral neurons (s-LNv, i.e., M cells) and PDF-negative dorsal lateral neurons (LNd) (plus at least fifth s-LNv, i.e., E cells), respectively.^52–55^ The M cells principally control the free-running rhythm in DD, and genetic ablation of these cells causes arrhythmicity. Furthermore, the activity time is correlated with the cellular period of E cells, whereas the period is correlated with that of M cells.^56^

Fly PDF has been suggested as the counterpart of mammalian VIP in the central clock function. Besides, the fly neuropeptide ion transport peptide (ITP), which is expressed in two neurons of E cells, may potentially be the counterpart of AVP.^55^ These analogous structures fit well with the dual oscillator model, predicting VIP and AVP neurons as M and E cells of the SCN, respectively. However, there are significant differences between the fly and mouse central circadian clocks that argue against such a simple analogy. For example, TTFLs in fly M cells are essential for pacemaking the free-running rhythm in DD.^54,55^ In contrast, manipulation of TTFL period or its disruption in VIP neurons has minimal effects on the behavioral rhythm, suggesting that TTFLs in VIP neurons may be dispensable for the ensemble pacesetting, although VIP peptide is critical.^13–16,18^ Instead, our previous results indicated that AVP neurons are the principal determinant of the free-running period in DD.^12,16,17^ In flies, Ca^2+^ rhythms in M and E cells show their peaks 3∼4 h before the M and E locomotor activities, respectively, tracking their respective behaviors.^57^ However, such temporal relationships have not been observed for VIP and AVP neurons.^11,16,21,27,28^ In particular, the in vivo Ca^2+^ activity of AVP neurons measured by fiber photometry is close to the sine curve with its peak in the early morning, whereas that of VIP neurons is close to the on and off square pulses, which almost delineate the (subjective) day under various LD conditions and even in a genetically modified free-running rhythm (*Avp-Bmal1^−/−^*mice).

Recently, CCK neurons of the SCN were found to be another candidate for the E cells because their Ca^2+^ rhythm tracks the E locomotor activity under various lighting conditions, although ablation of CCK neurons does not reduce E activity.^21^ Their ablation, but not a TeNT expression, shortens the free-running period in DD by 10∼20 min, suggesting that they are a part of the slow oscillators. Although these neurons play an important role in regulating circadian behavior under long-day photoperiod, the current study showed that they are a weak, unstable oscillator in the absence of VIP signaling. Therefore, at least in DD, CCK neurons are unlikely to be functional E cells.

TeNT expression in AVP neurons did not abolish E or M locomotor activities, but instead altered their phase relationship, resulting in an apparent diurnal behavior rhythm in most mice examined. This phenotype was consistent with the lengthened activity time in *Avp-Bmal1^−/−^*and *Avp-Vgat^−/−^* mice,^12,27^ suggesting that AVP neurons have two parallel functions: pacesetting of the ensemble period and phase setting of both E and M locomotor activities. This observation is another reason why we consider the simple projection of the fly M and E model onto the mammalian SCN network to be unlikely. In this context, we consider that the reduced M and E locomotor activities in VIP-deficient^6,8–10^ and *Avp-Bmal1^−/−^* mice^12^, respectively, do not reflect the attenuation of M and E oscillators, but result from the shorter and longer SCN ensemble periods.

To begin with, the central circadian clocks of mammals and flies have two major differences in their design.^54,55^ First, the fly clock network consists of seven discrete neuronal clusters that differ from other brain cells in that they have TTFLs, whereas in mammals, most SCN cells, and even most brain cells, have TTFLs. Second, most fly clock cells, including M and E cells, are sensitive to and directly reset by the external light, whereas the light is conveyed to the SCN core only via the retina. Therefore, it is not surprising that the logical structure of the central clock differs between mammals and flies. Having many clock neurons in the SCN that oscillate with widely differing phases, the phase distribution of the clock neurons, rather than simply two oscillators, may determine the onset and offset of locomotor activity in mammals.^55,58–60^

At the same time, we can still learn many things from the logic of the fly’s central clock network. For example, PDF signaling from M cells (s-LNv) delays the Ca^2+^ rhythm of E cells (LNd).^57,61^ In addition, PDF may entrain the intrinsically short-period LNd neurons by delaying the entrance of PER into the nucleus to accumulate PER protein and subsequently generate a more forceful negative feedback on s-LNv neurons to result in a greater amplitude of the central clock system.^62^ Such a scheme of delay and amplification by PDF signaling resembles our SNFL model that VIP signaling delays and amplifies AVP neuronal oscillation.^29–32^

In most species, the behavioral free-running periods differ significantly from 24 h and are adjusted to 24 h by the external LD cycle. This mechanism has been considered advantageous for stabilizing the phase relationship between the LD cycle and the behavior rhythm.^63^ A similar strategy may be used within the central clock network to stabilize the ensemble period, in which the VIP signal entrains AVP neuronal oscillators daily. Such a network architecture may also be adaptive to the photic entrainment.

## Acknowledgements

We thank H. Okamoto for the *Avp-Cre* mouse; S. Horike and T. Daikoku for the *Vip-tTA* mouse; Z. J. Huang and I. Fukunaga for the *Cck-ires-Cre* mouse; Z. J. Huang for the *Vip-ires-Cre* mouse; F. Zhang for the *Rosa26-LSL-SpCas9-2A-EGFP* mouse; C.J. Weitz for the *Bmal1^flox^* mouse; Penn Vector Core for *pAAV2-rh10*; D. Kim & GENIE Project for *pGP-AAV-CAG-FLEX-jGCaMP7s-WPRE* and *pAAV.CAG.Flex.NES-jRGECO1a.WPRE.SV40*; H. Kwon for *pAAV-TRE-EGFP* and *pAAV-TRE-ChR2-YFP*; *pAAV-TRE-ChrimsonR-mCherry* for A. Ting; *pAAV-flex-taCasp3-TEVp* for N. Shah and J. Wells; *pX333* for A. Ventura; *pAAV-EF1a-DIO-mCherry* for B. Roth; and *pAAV-EF1α-DIO-GFP::TeNT* for R.D. Palmiter; We thank all lab members, including M. Kawabata and Y. Nishiwaki.

## Author contributions

M.W. and M.M. designed research; M.W., Y.T., Y.P., A.M., T.M., and M.M. performed research; M.W., Y.T., and M.M. analyzed data; M.W. and M.M. wrote the paper.

## Material and methods

### Animals

*Avp-Cre* (C57BL/6J-*Tg(Avp-icre)#Meid*/Rbrc, RBRC12048) and *Vip-tTA* (B6(Cg)-*Vip^em^*^1^(tTA2)*^Miem^*/Rbrc, RBRC12109) mice were reported previously.^10,12^ *Cck-ires-Cre* (*Cck^tm^*^1^*^.^*^1^(cre)*^Zjh^*/J, JAX:012706),^22^ *Rosa26-LSL-SpCas9-2A-EGFP* (B6J.129(B6N)-*Gt(ROSA)26Sor^tm^*^1^(CAG–cas9*,–EGFP)*^Fezh^*/J, JAX:026175),^64^ *Vip-ires-Cre* (*Vip^tm^*^1^(cre)*^Zjh^*/J, JAX:010908),^22^ and *Bmal1 flox* (B6.129S4(Cg)-*Bmal1^tm1Weit^*/J, JAX:007668) mice were obtained from Jackson Laboratory.^65^ All lines were congenic on C57BL/6J. We compared the mutant mice with controls whose genetic backgrounds were comparable. *Avp-Cre*, *Cck-ires-Cre*, *Vip-ires-Cre*, and *Rosa26-LSL-SpCas9-2A-EGFP* mice were used in hemizygous or heterozygous condition. We used both male and female mice in our experiments. Whether we pooled data from both sexes or analyzed them separately, the conclusions we reached remained the same. The sex of individual mice is described in Supplemental Figures. Mice were maintained under a strict 12-h light/12-h dark cycle in a temperature- and humidity-controlled room and fed ad libitum. All experimental procedures involving animals were approved by the appropriate institutional animal care and use committees of Kanazawa University.

### Viral vector and surgery

The AAV-2 ITR containing plasmid *pGP-AAV-CAG-FLEX-jGCaMP7s-WPRE* (Addgene plasmid #104495, a gift from Dr. Douglas Kim and GENIE Project),^19^ was obtained from Addgene. *pAAV-TRE-ChR2-YFP* (Addgene #171622, a gift from Hyungbae Kwon),^66^ was modified to construct *pAAV-TRE-jRGECO1a* by replacing *ChR2-YFP* cDNA with a *jRGECO1a* cDNA fragment amplified by PCR from the plasmid *pAAV.CAG.Flex.NES-jRGECO1a.WPRE.SV40* (Addgene #100852, a gift from Douglas Kim & GENIE Project),^20^ using the following primers: 5’-CTAGGATCCGCCACCATGCTGCAGAACGAGCTTGC-3’ and 5’-CTGAAGCTTTCACTTCGCTGTCATCATTTGTAC-3’. Then, an EcoRI-HindIII fragment of the resultant plasmid containing *jRGECO1a* sequence was used to replace an EcoRI-HindIII fragment containing *ChrimsonR-mCherry* from *pAAV-TRE-ChrimsonR-mCherry* (Addgene #92207, a gift from Alice Ting).^67^ *pAAV-CAG-FLEX-taCasp3-TEVp* was made by replacing a BamHI-EcoRV fragment containing *EGFP* of *pAAV-CAG-FLEX-EGFP*^27^ with a BamHI-EcoRV fragment containing *taCasp3-TEVp* from *pAAV-flex-taCasp3-TEVp* (Addgene #45580, a gift from Nirao Shah & Jim Wells).^68^ *pAAV-U6-gVipr2-EF1α-DIO-mCherry*, a plasmid for CRISPR-Cas9-mediated *Vipr2* gene disruption, was generated as follows. The target sites for CRISPR-Cas9 were designed by CRISPOR (http://crispor.tefor.net/).^69^ Two sequences targeting *Vipr2* gene were selected: 5’-TGCTGGCGCCCGGCAGACGT-3’ and 5’-CCAGATTTCATAGATGCGTG-3’. Oligonucleotides encoding the guide sequences were cloned into the BbsI and BsaI sites of *pX333* (Addgene #64073, a gift from Dr. Andrea Ventura).^70^ Then, a fragment containing two tandem units of *U6-gRNA* was amplified by PCR, using the following primers: 5’-agtacgcgTCGAGCATGCTCGAGAATGG-3’ and 5’-agtacgcgtCGGGTACCCCATTTGTCTGC-3’, and cloned into the MluI site of *pAAV-EF1a-DIO-mCherry* (a gift from Dr. Bryan Roth) as described previously.^71^ *pAAV-U6-gControl-EF1α-DIO-mCherry* contains spacer sequences from *pX333* instead of gRNA sequences for *Vipr2*. *pAAV-EF1α-DIO-GFP::TeNT* was kindly provided by Dr. Richard D. Palmiter.^72^

Recombinant AAV vectors (AAV2-rh10) were produced using a triple-transfection, helper-free method and purified as described previously.^12^ The titers of recombinant AAV vectors were determined by quantitative PCR: AAV-*CAG-DIO-jGCaMP7s*, 3.4 × 10^13^; AAV-*TRE-jRGECO1a*, 8.6 × 10^12^; AAV-*CAG-FLEX-taCasp3-TEVp*, 5.7 × 10^12^; AAV-*U6-gVipr2-EF1α-DIO-mCherry*, 6.8 × 10^12^; AAV-*U6-gControl-EF1α-DIO-mCherry*, 2.6 × 10^12^; AAV-*EF1α-DIO-GFP::TeNT*, 1.8 × 10^12^ and AAV-*CAG-FLEX-EGFP*, 1.0 × 10^13^ genome copies/ml. Stereotaxic injection of AAV vectors was performed as described previously.^12^ Two weeks or immediately after surgery, we began monitoring the mice for their locomotor activity.

### In vivo fiber photometry

We used 8 *Avp-Cre; Vip^tTA/tTA^*, 6 *Cck^wt/ires-Cre^; Vip^tTA/tTA^*, 8 control (5 *Avp-Cre; Vip^wt/tTA^* and 3 *Cck^wt/ires-Cre^; Vip^wt/tTA^*), and 4 *Avp-Bmal1^−/−^* × *Vip-tTA* (*Avp-Cre; Bmal1^flox/flox^; Vip^wt/tTA^*) mice. Mice were anesthetized by administering a cocktail of medetomidine (0.3 mg/kg), midazolam (4 mg/kg), and butorphanol (5 mg/kg) and were secured to the stereotaxic apparatus (Muromachi Kikai). Lidocaine (1%) was applied for local anesthesia before making the surgical incision. We drilled a small hole in the exposed region of the skull using a dental drill. We injected 1.0 μL of the mixture of AAV-*CAG-FLEX-jGCaMP7s* and AAV-*TRE-jRGECO1a* (ratio 1:1, flow rate = 0.1 μL/min) into the right SCN (posterior: 0.5 mm, lateral: 0.25 mm, depth: 5.7 mm from the bregma) with a 33 G Hamilton syringe (1701RN Neuros Syringe, Hamilton) to label AVP and VIP neurons. We then placed an implantable optical fiber (400 μm core, N.A. 0.39, 6 mm, ferrule 2.5 mm, Thorlabs or RWD) above the SCN (posterior: 0.2 mm, lateral: 0.2 mm, depth: 5.4 mm from the bregma) with dental cement (Super-bond C&B, Sun Medical). The dental cement was painted black. Atipamezole (0.3 mg/kg) was administered postoperatively to shorten the anesthetized period. Mice were used for experiments 2 to 7 weeks after the virus injection and optical fiber implantation. Their ages ranged from 3 to 10 months, including both males and females.

A fiber photometry system (FP3002, Neurophotometrics) was used to record the calcium signal of SCN neurons in freely moving mice.^73,74^ Excitation light sources were a 470-nm LED for detecting Ca^2+^-dependent jGCaMP7s fluorescence signal (F470), a 560-nm LED for detecting Ca^2+^-dependent jRGECO1a fluorescence signal (F560), and a 415-nm LED for Ca^2+^-independent isosbestic fluorescence signal (F415). The duration of the excitation lights was 50 ms, and the onsets of the excitation timing of the LEDs were interleaved. The lights passed through excitation bandpass filters, dichroic mirrors, and then to the animal via fiber-optic patch cords (BBP(4)_400/440/900-0.37_1m_FCM-4xFCM_LAF, MFP_400/440/LWMJ-0.37_1m_FCM-ZF2.5_LAF, Doric Lenses) and the implanted optical fiber. Subsequently, all signals were detected using a CMOS camera through the optical fibers, dichroic mirrors, and emission bandpass filters. The green emission signal and the red emission signal were recorded separately and simultaneously. Ca^2+^ signals were recorded every 10 min for 30 s using Bonsai software at a sampling rate of 6.6 Hz for each color. The excitation intensities of the 470-nm, 560-nm, and 415-nm LEDs at the tip of the patch cord on the animal side ranged from 60 μw to 110 μw. During recording, the mouse was housed in a 12-h light–dark cycle for more than 7 days (LD condition) and then transferred to continuous darkness for more than 21 days (DD condition) in a custom-made acrylic cage surrounded by a sound-attenuating chamber.

Ratio (R) was defined as the ratio between F470 and F415 in green channel (F470/F415 green) or the ratio between F560 and F415 in red channel (F560/F415 red) for calibration and reducing motion artifacts. R was averaged within a 30-s session.^27^ To detrend the gradual decrease of the signal during the recording days, ± 12 h average from the time (145 points) was calculated as baseline (R_0_). The data were subsequently detrended by the subtraction of R_0_ (ΔR). Then, the ΔR/R_0_ value was calculated. To determine the daily peak phase of jGCaMP7s or jRGECO1a Ca^2+^ signal, ΔR/R_0_ were smoothened with a 21-point moving average, then the middle of the time points crossing value 0 upward (Ca^2+^ onset) and downward (Ca^2+^ offset) were defined as the midpoint that reflect the peak phases. Additionally, the intervals between two adjacent midpoints were defined as the daily periods (Figure 1E).^12^ Because VIP-Ca^2+^ rhythm in VIP deficient mice attenuated drastically in DD, we failed to determine its Ca^2+^ onset, offset, and daily period at the late stage. A double-plotted actogram of the jGCaMP7s or jRGECO1a signal was prepared by converting all ΔR/R_0_ to positive values by subtracting the minimum value of ΔR/R_0_. Subsequently, these values were multiplied by 100 or 1,000 and rounded off. The plots were made via ClockLab (Actimetrics) with normalization in each row to emphasize weak VIP-Ca^2+^ and CCK-Ca^2+^ rhythmicity in VIP-deficient mice at the late stage in DD. To quantitively analyze the weak rhythms and their changes, periods and amplitudes of Ca^2+^ signals (ΔR/R_0_) were calculated by periodogram with 10-min bins for all three weeks and shifting five-day time windows in DD, respectively (Figures 2, 3, S1, S2 and S3).

During the fiber photometry recordings, the animal’s locomotor activity was monitored using an infrared sensor (AMN31112 and Arduino) in 10-min bins. A double-plotted actogram of locomotor activity was also prepared and overlaid on that of the jGCaMP7s and jRGECO1a signal. The onset and offset of locomotor activity were determined using the actograms. Initially, we attempted to automatically detect the onset and offset; however, it was followed by a manual visual inspection and modifications by the experimenter. The intervals between two adjacent locomotor onsets were defined as the daily periods (Figure 1, 5F). Because VIP-deficient mice did not show clear morning locomotor activity, we did not determine the locomotor activity offset for these mice. To calculate CT of the midpoint phases of jGCaMP7s and jRGECO1a signals, we have defined the regression line of locomotor activity onsets as CT12. Period and amplitude of locomotor activity were also calculated by χ^2^ periodogram analyses for all days or 5 consecutive days in DD to analyze weak rhythmicity of VIP-deficient mice (Figures 2, 3, S1, S2 and S3).

We confirmed the jGCaMP7s and jRGECO1a expressions and the position of the optical fiber by slicing the brains into 30 μm coronal sections using a cryostat (Leica). The sections were mounted on glass slides with a mounting medium (VECTASHIELD HardSet with DAPI, H-1500, Vector Laboratories; Dako Fluorescence Mounting Medium, Agilent Technologies) and observed via epifluorescence microscope (BZ-9000, Keyence).

### Behavioral analyses

Male and female *Vip-Casp3* (*Vip^wt/ires-Cre^* mice injected with AAV-*CAG-FLEX-taCasp3-TEVp*) (Figure S4), *Avp-Vipr2^−/−^* (*Avp-Cre; Rosa26-LSL-Cas9-2A-EGFP* mice injected with AAV-*U6-gVipr2-EF1α-DIO-mCherry*) or control (*Avp-Cre; Rosa26-LSL-Cas9-2A-EGFP* mice injected with AAV-*U6-gControl-EF1a-DIO-mCherry*) (Figure 4), and *Avp-TeNT* (*Avp-Cre* mice injected with *AAV-EF1α-DIO-GFP::TeNT*) or control (*Avp-Cre* mice injected with AAV-*CAG-FLEX-EGFP*) (Figure 6) mice, aged 8 to 30 weeks, were housed individually in a cage placed in a light-tight chamber (light intensity was approximately 100 lux). Spontaneous locomotor activity (home-cage activity) was monitored by infrared motion sensors (O’Hara) in 1-min bins with a custom-made program as described previously.^12^ Actogram, activity profile, and χ^2^ periodogram analyses were performed via ClockLab (Actimetrics). The free-running period and amplitude were measured by periodogram for all days in DD (Figure 6) or the last 2 weeks in DD (Figure S4). In the case of *Avp-Vipr2^−/−^* and control mice, their periods and amplitudes were measured by periodogram for 10 days in 1st or 2nd stage in DD (Figure 4).

### Immunohistochemistry

Immunostaining was performed as described previously.^10,12,71^ Mice were killed approximately at ZT2 by transcardial perfusion of PBS followed by 4% paraformaldehyde (PFA) in PBS. Then, the mice brains were postfixed in the 4% PFA at 4°C overnight, followed by immersion in 30% sucrose solution at 4°C for 2 days. Serial coronal brain sections (30 μm thickness) were made with a cryostat (CM1860, Leica) and collected in 4 series—one of which was further immunostained. For immunofluorescence staining, sections were washed with PBS containing 0.3% Triton X-100 (PBST) and blocked with PBST plus 3% BSA (blocking solution). Then, slices were incubated overnight with the designated primary antibodies in the blocking solution at 4°C. Antibodies used were rabbit anti-GFP antibody (1:1,000; A11122, Thermo Fisher Scientific) and rabbit anti-VIP antibody (1:1,000; #20077, Immunostar). Then, slices were washed with PBST, followed by incubation with the designated secondary antibodies in blocking solution for 4 h. Secondary antibodies used in this study were Alexa Fluor 488–conjugated donkey anti-rabbit IgG antibody (1:2,000, A-21206; Thermo Fisher Scientific) and Alexa Fluor 594–conjugated donkey anti-rabbit IgG antibody (1:2,000, A-21207; Thermo Fisher Scientific). After incubation with secondary antibody, slices were washed with PBS, mounted on slide glasses, air dried, and coverslipped using Mounting Medium (H-1500, Vector Laboratories; Dako Fluorescence Mounting Medium, Agilent Technologies). Images were taken using an epifluorescence microscope (BZ-9000, Keyence).

### Statistical analysis

All results are expressed as mean ± SEM. Statistical analyses were performed using Prism 8.0 software (GraphPad). For comparisons of 2 groups, two-tailed Student’s t tests were performed. For comparisons of multiple groups with no difference of variance by Bartlett test, two-way repeated measures ANOVA followed by post hoc Ryan test were performed. For circular data, Rayleigh test, Watson–Williams test, and Harrison–Kanji test were performed with Circstat MATLAB Toolbox for Circular Statistics.^75^ All P values less than 0.05 were considered as statistically significant. Only relevant information from the statistical analysis was indicated in the text and figures.

